# Exploitation of CD3ζ to enhance TCR expression levels and antigen-specific T cell function

**DOI:** 10.1101/2024.02.14.580308

**Authors:** Abdullah Degirmencay, Sharyn Thomas, Angelika Holler, Hans J Stauss

## Abstract

The expression levels of TCRs on the surface of human T cells define the avidity of TCR-HLA/peptide interactions. In this study, we have explored which components of the TCR-CD3 complex are involved in determining the surface expression levels of TCRs in primary human T cells. The results show that there is a surplus of endogenous TCR α/β chains that can be mobilised by providing T cells with additional CD3γ,δ,ε,ζ chains, which leads to a 5-fold increase in TCR α/β surface expression. The analysis of individual CD3 chains revealed that provision of additional ζ chain alone was sufficient to achieve a 3-fold increase in endogenous TCR expression. Similarly, CD3ζ also limits the expression levels of exogenous TCRs transduced into primary human T cells. Interestingly, transduction with TCR plus CD3ζ not only increased surface expression of the introduced TCR, but it also reduced mispairing with endogenous TCR chains, resulting in improved antigen-specific function. TCR reconstitution experiments in HEK293 cells that do not express endogenous TCR or CD3 showed that TCRα/β and all four CD3 chains were required for optimal surface expression, while in the absence of CD3ζ the TCR expression was reduced by 50%. Together, the data show that CD3ζ is a key regulator of TCR expression levels in human T cells, and that gene transfer of exogenous TCR plus CD3ζ improved TCR surface expression, reduced TCR mispairing and increased antigen-specific function.

## INTRODUCTION

T cells play a crucial role in the adaptive immune system, recognising a vast array of antigens through specialised T cell receptors (TCRs), consisting of αβ or γδ chains(1). Produced in the endoplasmic reticulum (ER) of T cells, TCR chains assemble with four CD3 chains, which form εδ and εγ heterodimers and ζζ homodimers (2). It has been shown that the TCRα chain pairs with the CD3 εδ heterodimer, and TCRβ pairs with the εγ heterodimer. The fully assembled TCR-CD3 complex is formed after ζζ homodimers join this pentameric structure through TCRα chain binding(2). The assembly of the TCR-CD3 complex is vital for TCR surface expression since in the absence of CD3 the TCR α and β chains undergo rapid degradation in the ER(3). In addition to enabling TCR surface expression, CD3 plays a key role in transmitting T cell activation signals through ITAMs found in the cytoplasmic tails of each CD3 chain(4).

Being a key component of TCR surface expression, CD3 is also crucial in TCR gene therapy, where efficient TCR-CD3 assembly is required for optimal antigen-specific function. We have previously shown that CD3 limits the TCR surface expression of TCR gene engineered murine T cells. We found that the transduction of CD3γ,δ,ε,ζ together with TCRα/β markedly enhanced TCR expression levels of engineered T cells and improved their ability to protect against tumour growth in vivo(5). However, it has remained unclear whether all four CD3 chains are required to achieve enhanced TCR expression, and whether the findings in mouse T cells would also apply to humans.

In this study we have performed a detailed analysis of the role of TCRα/β and CD3γ,δ,ε,ζ in the regulation of TCR surface expression in primary human T cells. We found that TCR surface levels are primarily regulated by CD3ζ, and that an abundant pool of endogenous TCRα/β and CD3γ,δ,ε chains can be recruited for increased surface expression by providing human T cells with additional CD3ζ. In the setting of TCR gene therapy, the co-transduction of TCR and CD3ζ increased the surface expression of the introduced TCR and enhanced the antigen-specific function.

## METHODS

### Cell culturing

Human PBMCs (Human Peripheral blood mononuclear cells) and HLA-A2+ T2 cells (can be efficiently loaded with exogenous peptides as they lack the transporter associated with antigen processing) were cultured in RPMI 1640 media (Lonza) supplemented with 10% FCS (Merck), 2mM L-Glutamine (Gibco) and 100U/ml Penicillin/Streptomycin (Invitrogen). HEK293T (Human Embryonic Kidney Epithelial) packaging cells were cultured in IMDM media (Lonza) supplemented with 10% FCS and 2mM L-Glutamine and 100U/ml Penicillin/Streptomycin.

### Activation of human PBMCs and CD8+ T cells

Human PBMCs were obtained from healthy volunteers via National Health Blood Transfusion Service (Approved by UCL Research Ethics Committee, Project ID: 15887/001). 48 hours prior to retroviral transduction, bulk PBMCs were activated at 1×10^6^ cells/ml with 20ml/ml anti-CD3/CD28 dynabeads (Gibco) and 30U/ml Roche IL-2.

For T cell functional assays, PBMCs were MACS (Magnetic-Activated Cell Sorting) sorted for CD8+ T cells following the manufacturer’s instructions (Miltenyi Biotech). Sorted CD8+ T cells were activated at 1×10^6^ cells/ml with 20ml/ml anti-CD3/CD28 dynabeads and 30U/ml Roche IL-2.

### Antibodies and peptides

The following antibodies were used for the flow cytometric analysis of the cells: anti-human antibodies CD3-PE-Cy7 (SK7, BioLegend), TCRα/β-PE (IP26; Invitrogen), IgG1-PE (Invitrogen), IL-2 APC (MQ1-17H12, eBioscience) and IFN-g-PE (B27; Invitrogen). Other antibodies used were anti-murine CD19-eFluor450 (1D3; Invitrogen), V5-APC (rabbit polyclonal; abcam) and purified myc (Bio-Rad). Live/Dead-eFluor780 (Invitrogen) was used to identify live cells. Peptides used for the T cell functional assays were: pCMVpp65 (NLVPMVATV) for the CMV TCR and pHA1 (VLHDDLLEA) for the HA-1.m2 and HA-1.m7 TCRs. The pHA2 (YIGEVLVSV) peptide was used as a control peptide.

### Retroviral vectors

DNA constructs were cloned into retroviral pMP71 vectors. For the CD3 constructs, IRES GFP was placed at the 3’ end and viral 2A sequences were used to separated constructs with polycistronic genes. The TCR constructs consisted of a TCRα chain, a viral P2A sequence, a TCRβ chain, a viral T2A sequence, and truncated murine CD19. A V5 tag was incorporated upstream of the TCR α variable domain and two myc tags were placed upstream of the TCR β variable domain.

### Retrovirus production and cell transductions

Retrovirus production and the transduction of primary human T cells were performed as previously described(6). For the retroviral co-transduction of HEK293 cells, 2×10^5^ HEK293 cells were seeded into wells of a 24 well tissue culture treated plate in 500ul IMDM media and incubated for 30 mins at 37°C. After ensuring the cells were attached, media was discarded carefully and 250ml of each of the stated CD3 and TCR viral supernatants were added. Transduction was performed by centrifugation at 32°C, 2000rpm, 90 mins, no brake. The viral supernatant was discarded, and each well was supplied with 2ml fresh IMDM. 48-72h post transduction, cells were stained for Live/dead, and the antibodies stated in the text. Data was collected by LSRFortessa (BD Biosciences) and the analysis was done by FlowJo software.

### Intracellular Cytokine Staining

3×10^5^ T2 cells were loaded with the stated concentrations of relevant or irrelevant peptide for 2 hours at 37°C, washedand thenco-cultured with 7-10 days post-transduced 3×10^5^ CD8+ T cells for 18 hours. Assays were conducted in a 96 well plate, round bottom in a volume of 250 ul / well of RPMI media supplemented with Brefeldin A (Merck) at 1 ug/ul. Cells were stained for surface markers and washed. Fixation and permeabilization of the cells was performed by BD Cytofix/Cytoperm Kit as per manufactuer’s instructions. and then stained for IL-2 and IFN-γ for 1 hour at 4°C. Data was acquired on a LSRFortessa and analysed using FlowJo software.

## RESULTS

### 1. Endogenous CD3 levels limit TCR expression in human T cells

We designed a retroviral vector cassette encoding all four human CD3 chains separated by 2A sequences, and GFP separated by an IRES element (Fig 1A). We also produced vectors that contained only one CD3 chain (ζ or ε or δ or γ) and a control vector containing IRES GFP but no CD3 components. GFP expression was used to demonstrate successful transduction of primary human T cells, and TCR surface expression was determined in gated cells expressing similar levels of GFP (Fig 1b, c). Using antibodies specific for human α/β TCR and human CD3ε we found that the expression levels of TCR in T cells transduced with the vector encoding all CD3 chains was approximately 5-fold higher than in T cells transduced with the GFP control vector (Fig 1c). The transduction of individual CD3 chains revealed that ζ was able to increase TCR expression by nearly 3-fold, while transduction with ε resulted in only modest improvement of TCR expression. Surprisingly, provision of additional γ or δ significantly decreased TCRα/β surface expression in human T cells (Fig 1d). This is most likely due to a competition for CD3ε whereby high levels of γ leads to high levels of γ/ε dimers and a reduction in δ/ε dimers, while high levels of δ reduces γ/ε dimer formation. The reduction of either δ/ε dimers or γ/ε dimer impairs the assembly of and surface expression of intact TCR/CD3 complexes in T cells with excess γ or δ, respectively. The observed changes in TCRα/β surfaces expression were mirrored by similar changes in CD3ε, although the intensity of CD3ε staining was less than the TCR staining.

**Figure 1.**
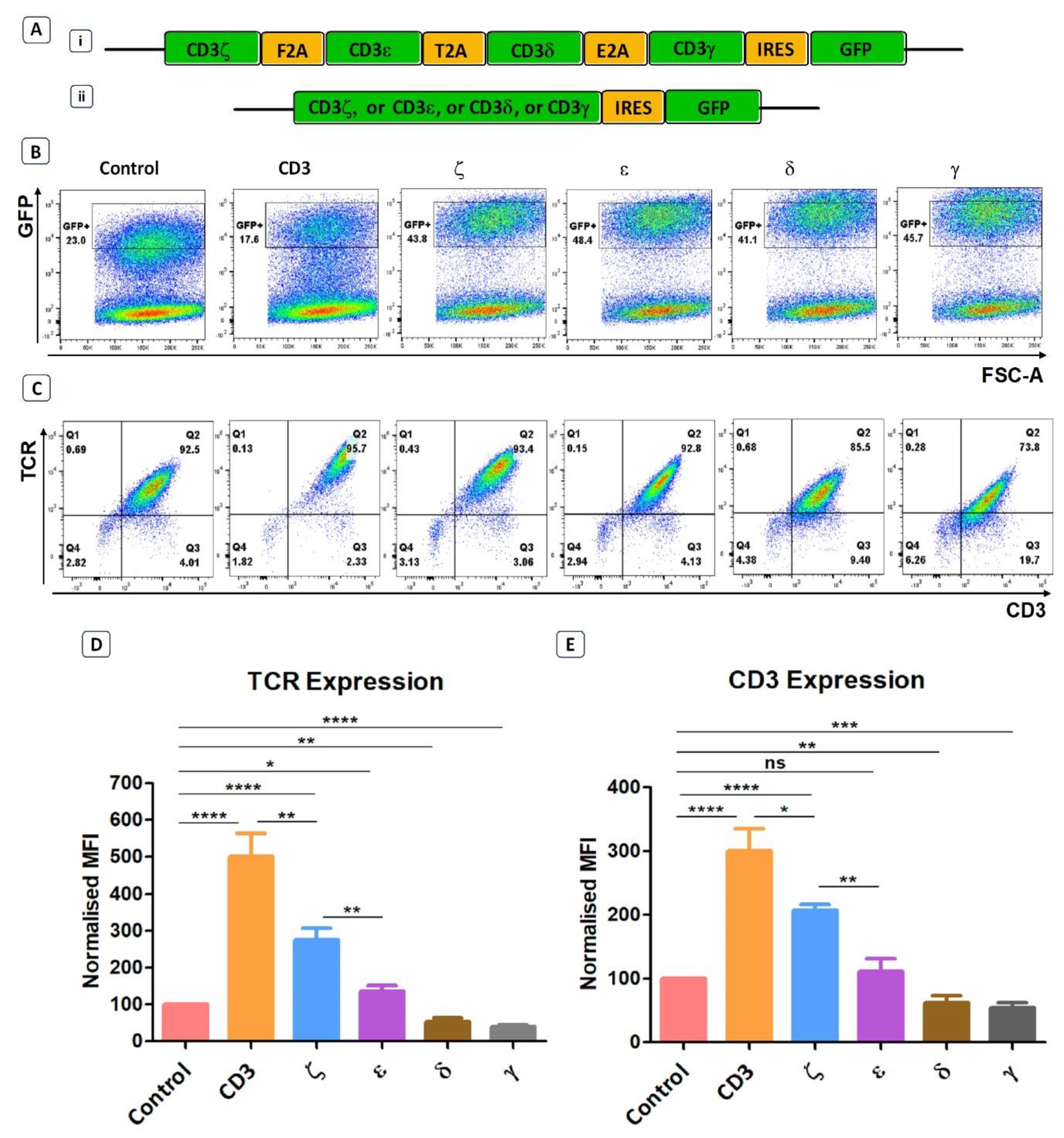
Endogenous CD3 limits TCR expression in human T cells. **a)** Schematic representations of the CD3 retroviral vectors encoding all 4 CD3 chains (i) or only one of the CD3 chains (ii) and used to transduce primary T cells. **b)** Representative example of seven independent experiments of primary T cells transduced with either a GFP control construct, or the construct containing all CD3 genes, or individual genes. Shown is GFP expression, indicating transduction efficiency. **c)** Representative plots of seven independent experiments showing TCR and CD3 expression in GFP+ population defined by the gates in panel b). **d-e)** Pooled data bar graphs showing MFI (median fluorescence intensity) of TCR and CD3 expressions in GFP+ gated primary T cells transduced with control or CD3 constructs (n=7). MFI (Mean +/-SEM) data has been normalised to control. Unpaired t-test was applied; * p<0.05, ** p<0.01, *** p<0.001, **** p<0,0001, ns p>0.05.

### 2. Not all four CD3 chains are required to achieve optimal TCR surface expression

Next, we tested whether all CD3 chains were required for optimal expression increase of endogenous TCRs in primary human T cells. We first generated 4 vectors that contained three CD3 chains but lacked either γ,δ,ε or ζ. Transduction of T cells revealed that the lack of ζ almost completely abolished the ability of the introduced CD3 chains to increase TCR surface expression (Fig 2a, b). In contrast, the lack of γ or δ did not significantly reduce the level of TCR upregulation that is seen when all four CD3 chains were transduced into T cells, whereas significantly less TCR expression was observed when ε was the missing in the transduction vector (Fig 2a, b). This suggested that the endogenous levels of CD3ζ are the most rate limiting CD3 chain in determining TCR surface expression, followed by CD3ε, while endogenous CD3γ and δ are relatively abundant and do not limit TCR/CD3 assembly and surface expression. This interpretation is compatible with the observation that the transduction of T cells with a vector encoding only ζ,ε and lacking γ,δ resulted in substantial upregulation of TCR surface expression (Fig 2a, b).

**Figure 2.**
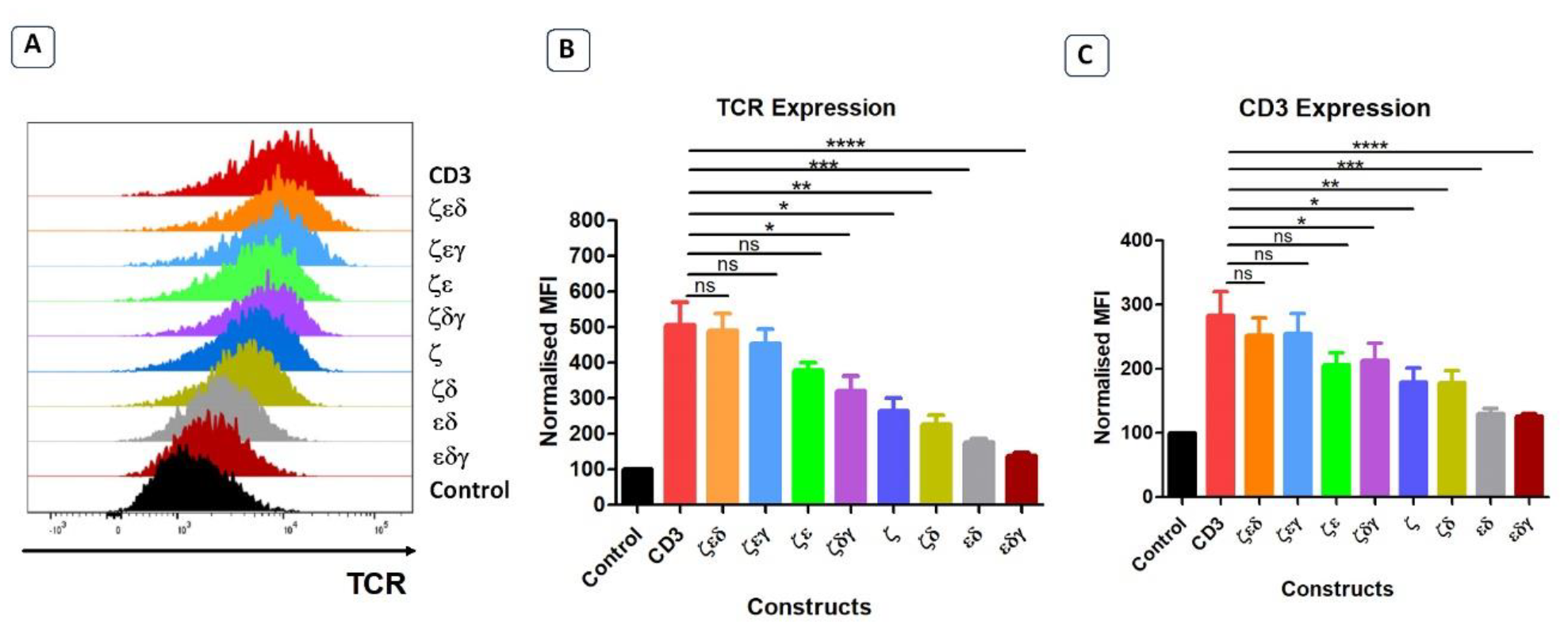
Not all four CD3 chains are required for maximal TCR expression. **a)** Representative histogram plot of five independent experiments showing the TCR expression in primary T cells transduced with control or stated CD3 constructs. Pooled data bar graphs showing the TCR (b) and CD3 (c) expressions of gated GFP+ T cells. MFI data (mean +/-SEM) has been normalised to control (n=5). Unpaired t-test was applied: * p<0.05, **, p<0.01, ***, p<0.001, ****, p<0.0001, ns p>0.05.

The surface expression profiles of CD3ε showed a similar pattern as the expression profiles of TCRα/β (Fig 2c). Interestingly, transduction of T cells with the vector containing only γ,δ,ζ resulted in a 2-fold increase in CD3ε surface expression, indicating that surplus endogenous ε chains were available for the assembly of additional TCR-CD3 complexes.

### 3. CD3 δ and ε are essential to reconstitute TCR surface expression in HEK293 cells

The previous experiments have identified which CD3 chains are rate limiting and which are abundant in primary human T cells. In the next set of experiments, we wanted to determine which CD3 chains are essential to achieve TCR surface expression. For this purpose, we used the HEK293T (Human Embryonic Kidney) cells that lack endogenous CD3 and TCR. We used vectors expressing all four CD3 chains, or various chain combinations as illustrated and described in figure 3a, c. We also used a vector encoding an HLA-A0201-restricted TCR specific for CMV, where the α and β chains were tagged with V5 and myc epitopes, respectively, to measure TCR expression with antibodies specific for these tags(7) (Fig 3b). Co-transduced HEK293T cells were analysed for GFP and CD19 expression, to identify GFP+CD19+ positive cells that were successfully co-transduced with both the CD3 and the TCR vectors (Fig 3c). Co-transduction of all four CD3 chains and the TCR resulted in efficient TCR α/β surface expression, whereas co-transduction of GFP control vector and TCR did not result in TCR surface expression (Fig 3c). We then tested vectors with three CD3 chains and found that the absence of the δ or ε chain nearly abolished TCRα/β surface expression (Fig 3c, d, f). In contrast, the effect of absence of the γ or ζ chain was less detrimental as TCR expression was retained, although much reduced compared to the expression achieved with all four CD3 chains (Fig 3c, d, f). We observed that the CD3 δ and ε chains were sufficient to achieve low levels of TCRα/β expression on the surface of transduced cells (Fig3c, d, f). As before, the expression pattern of CD3ε in the co-transduced HEK293T cells mirrored the observed TCRα/β expression (Fig 3d, e). These data suggest that CD3 δ/ε dimers are particularly important in initiating the assembly of a TCR-CD3 complex, which can then associate with γ/ε dimers and ζ/ζ homodimers for efficient migrating from the ER to the cell surface. In the absence of γ/ε and ζ/ζ some assembled TCRα/β-CD3δ/ε complexes seem to escape ER degradation and reach the cell surface, although at much lower levels than fully assembled TCR-CD3 complexes containing all CD3 chains. This particular importance of CD3δ/ε dimers in the TCR-CD3 complex is consistent with recent publications uncovering the detailed structure of the fully assembled TCRα/β-CD3γ,δ,ε,ζ complex(2,8,9)

**Figure 3.**
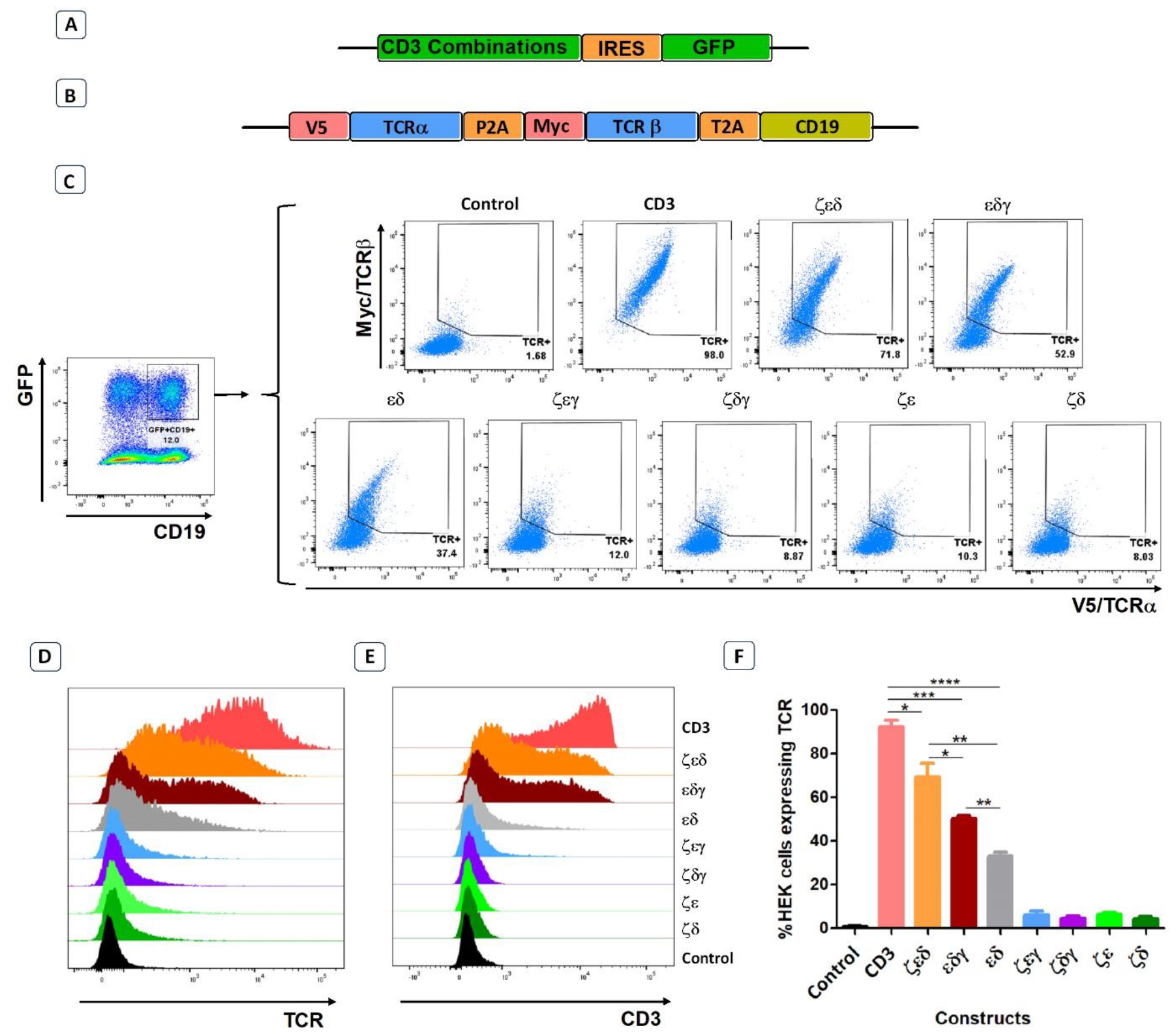
CD3εδ initiate TCR reconstitution in HEK293 cells. **a)** Schematic representation of the CD3 constructs retroviral vectors. GFP was used as a transduction marker. **b)** Schematic representation of the retroviral vector map of the CMV TCR construct. V5 and Myc tags allow the expression levels of TCRα and TCRβ chains to be determined through anti-V5 and anti-Myc antibodies, respectively. CD19 was used as a transduction marker. **c)** Representative example of three independent experiments showing the TCRα and TCRβ expressions of HEK293 cells co-transduced with control or CD3 constructs along with CMV TCR (GFP+CD19+ cells; left hand-side plot). **d-e)** Representative histogram plots of three independent experiments showing TCR **(d)** and CD3 **(e)** expressions of HEK293 cells co-transduced with control or CD3 constructs along with CMV TCR. **f)** Pooled data bar graphs showing the percentage of GFP+CD19+ gated HEK293 cells expressing the introduced CMV TCR (n=3). Unpaired t-test was applied; * p<0.05, **p<0.01, *** p<0.001, **** p<0,0001.

### 4. CD3ζ provision can be harnessed in TCR gene therapy

As primary human T cells contained surplus CD3γ, δ, ε chains, we explored whether the provision of additional ζ chains in the context of TCR gene therapy would be sufficient to enhance the expression levels and the antigen-specific function of introduced TCRs. Hence, we placed the human CD3ζ gene downstream of three different TCRs specific for the minor histocompatibility antigens HA-1.m2 and HA-1.m7, and for the pp65 antigen of CMV (Fig 4a). The TCR genes were tagged with the V5 and myc epitopes to measure TCRα and β expression, and CD19 was used to identify transduced T cells. For each TCR we also tested 3 variants that contained defined amino acid changes in the framework of the TCR variable domains that we previously demonstrated to improve TCR expression levels (6,7).

**Figure 4.**
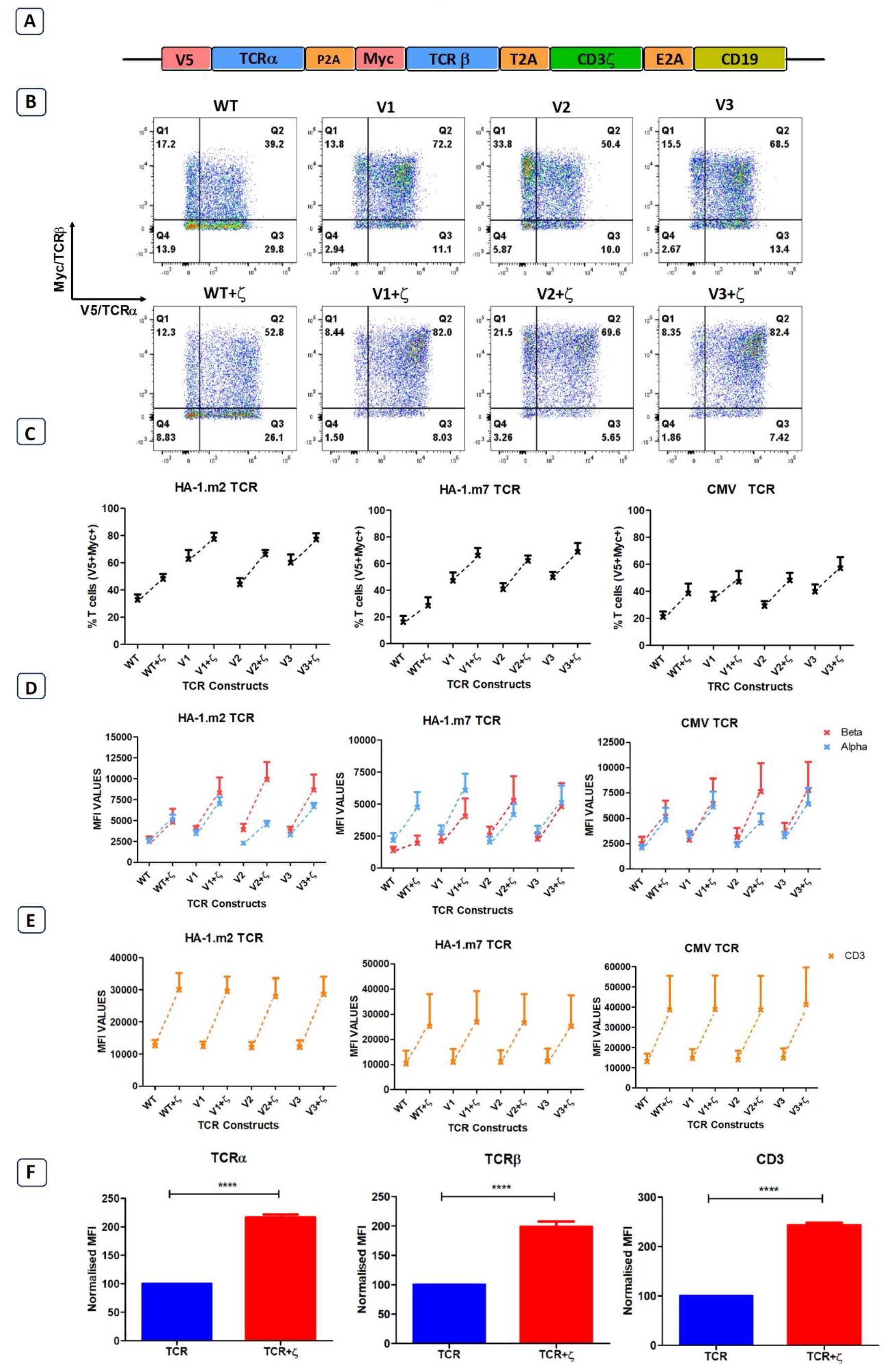
Following antigen-specific TCR transduction, CD3ζ provision enhances TCR expression and reduces mispairing. **a)** Schematic representation of the retroviral TCR+ζ constructs. TCR only vectors did not contain CD3ζ. **b)** Representative FACS plots of three independent experiments showing primary T cells transduced with the wild-type HA-1.m2 TCR (WT) or three variants V1, V2, V3 (top panel) and their CD3ζ added versions (bottom panel). Q2 numbers are the percentages of cells expressing both α and β chain of the introduced TCR. **c)** Pooled data (mean +/-SEM) of three independent experiments showing the percentage of transduced cells expressing both introduced TCR chains for all TCR versions tested with or without CD3ζ. **d-e)** Pooled data (mean +/-SEM) of three independent experiments showing TCRα/β **(d)** and CD3 **(e)** expressions of primary T cells transduced with the TCR versions of 3 antigen specific TCRs with or without CD3ζ. **f)** Summary pooled data of relative MFI of TCRα/β and CD3 expression of all 12 TCRs tested in 3 independent experiments, comparing transduction with TCR only, and TCR+ζ. The MFI values of TCR+ζ are relative to that of TCR only. Unpaired t-test was applied. **** p<0,0001.

Figure 4b shows a representative experiment of human T cells transduced with the HA1.m2 wild type TCR or with the three variants V1-3 (top panels). The bottom panels show expression levels of each TCR in the presence of additional CD3ζ. The data show that for each TCR, the inclusion of CD3ζ increased the percentage of T cells that express both introduced TCR chains (V5+myc+) and reduced the percentage of cells that are single positive for V5 or myc. In these single positive T cells the introduced V5-tagged α chains have mispaired with endogenous β chains, and the myc-tagged β chains have mispaired with endogenous α chains. Figure 4c shows that additional ζ consistently increased the percentage of T cells expressing the introduced α and β chains for all 12 TCRs tested. In addition to the increase in T cell numbers expressing both TCR chains, exogenous CD3ζ also increased the level of surface expression in transduced cells (Fig 4d), which correlated with increased expression levels of CD3ε (Fig 4e). Figure 4f shows the summary of TCRα, β and CD3ε expression levels of all 12 TCRs in the absence and presence of CD3ζ.

### 5. Additional CD3ζ boosted antigen specific cytokine expression in human T cells

As additional CD3ζ augmented the density of the introduced TCRs on the surface of human T cells, we tested whether this improved antigen specific T cell function. CD8+ T cells expressing either TCR-only, or the TCR+ζ combination of the HA-1.m2, HA-1.m7 or CMV-specific TCR were challenged with titrated concentrations of relevant peptide. In these functional experiments we compared the wild type TCR with the variant 3 modifications in the framework of the V-regions, because we previously showed that the V3 modifications were most effective in increasing TCR expression and function(10). This allowed us to test whether addition of CD3ζ can further improve the functionality of the V3 TCRs. Figure 5a shows that the addition of CD3ζ improved the antigen-specific IFN-γ and IL-2 production of the wild type HA1.m7 TCR and also of the V3 variant. The combination of V3+CD3ζ was most efficient in enhancing the poor antigen-specific response of the wild type HA1.m7 TCR. The analysis of the HA1.m2 and CMV TCRs confirmed that the V3 modification combined with CD3ζ was most effective in enhancing antigen-specific functionality of wild type TCRs, although the CMV-V3 TCR mounted a strong response in the absence of additional CD3ζ(Fig 5b).

**Figure 5.**
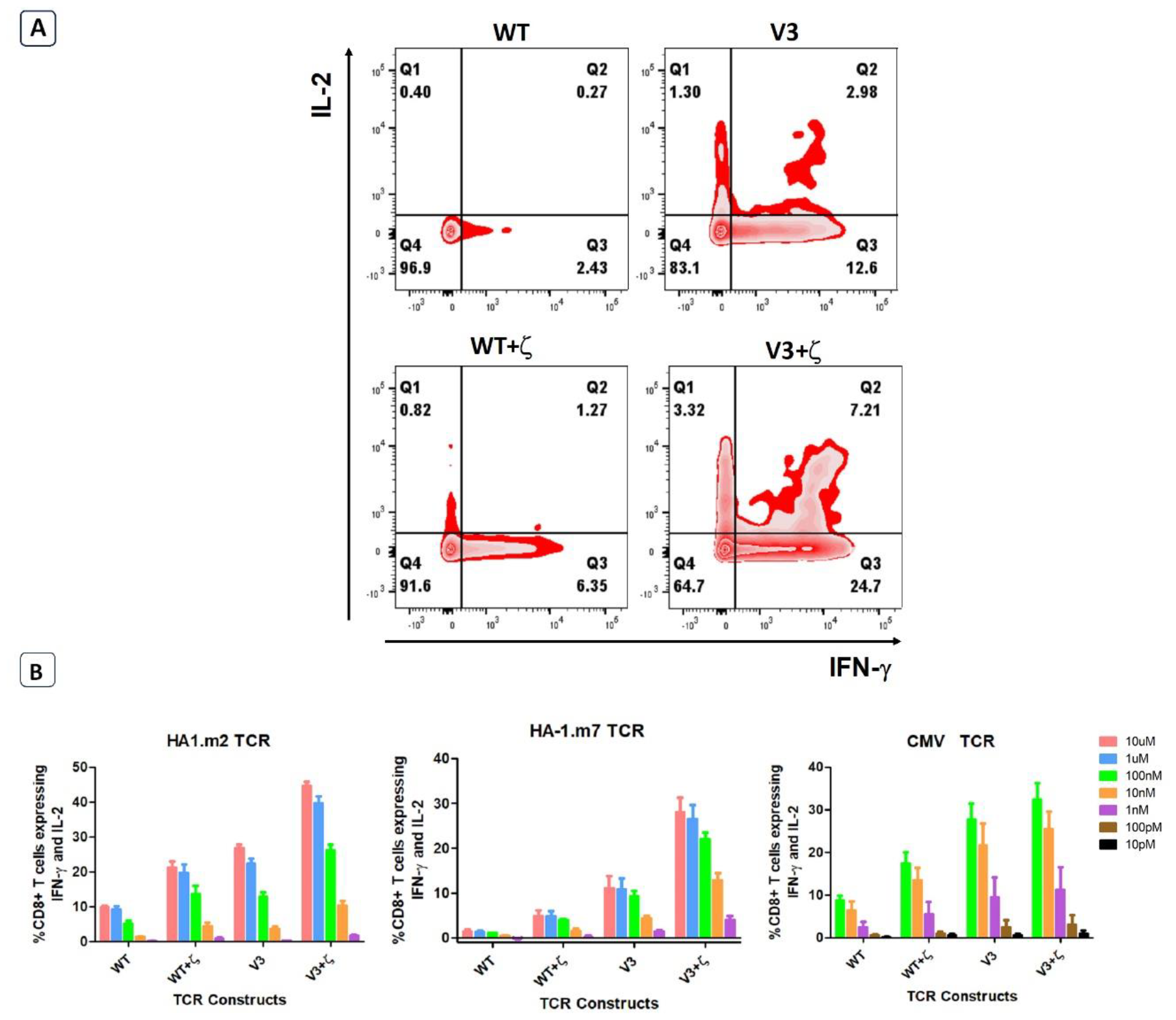
CD3ζ provision increases antigen specific cytokine production. 7-10 days post-transduced T cells were co-cultured with relevant or irrelevant peptide loaded T2 cells at 1:1 ratio. **a)** Representative FACS plots of three independent experiments showing the IL-2 and IFN-γ expression of primary T cells transduced with HA-1.m7 WT or V3 without CD3ζ (top panel) or with CD3ζ (bottom panel) following stimulation with 10uM peptide. **b)** Pooled data (n=3) showing the percentage of T cells expressing both IL-2 and IFN-γ at the stated peptide concentrations.

## DISCUSSION

In this study we showed that the level of TCR expression on the surface of human T cells is regulated by the CD3 chains. The majority of endogenously produced TCR α and β chains are not assembled into TCR-CD3 complexes and thus not expressed on the cell surface. Providing human T cells with all four additional CD3 chains resulted in a 5-fold increase in TCRα/β surface expression, suggesting that in physiologic conditions T cells utilise no more than 20% of the endogenously produced TCRα/β chains. We also found a hierarchy in the abundance of CD3 chains and identified γ as the most abundant chain, followed by δ, ε and ζ. Our data indicated that the ζ chain is the key regulator of TCR surface levels, since provision of additional εδγ had little effect on surface expression, while provision of additional ζ alone resulted in 3-fold increase of TCR on the cell surface. A critical role of CD3ζ in TCR expression and T cell function is supported by the observation that decreased ζ expression, associated with reduced TCR expression and impaired T cell function, has been described in tumour infiltrating T cells in various human cancers(11). Although several mechanisms may contribute to decrease ζ expression in tumour infiltrating T cells(12– 16) the restoration of sufficient CD3ζ expression levels provides an attractive target for cancer immunotherapy.

While CD3ζ determines TCR expression levels in primary human T cells, our experiments in HEK293T cells have shown that TCRα/β surface expression can be achieved in the absence of ζ. In contrast, the CD3δ and ε chains were strictly required to achieve any detectable TCRα/β expression in transduced HEK293T cells. This is consistent with recent structural details of the TCR/CD3 complex, showing a key role of the interaction between δ/ε dimer and the TCRα chain in enabling the assembly of the complete TCR-CD3 complex(8,9). Although CD3δ/ε was sufficient to achieve detectable TCR expression, it was very inefficient as the TCR surface levels were only 1/10 (check MFI) of what was achieved when all four CD3 chains were present. This suggests that, in the absence of γ and ζ, the majority of TCRα/β-CD3δ/ε complexes did not arrive at the cell surface and were instead degraded intracellularly. Interestingly, the addition of ζ to the TCRα/β-CD3δ/ε increased TCR surface expression levels to 80% (check MFIs) of the fully assembled TCR-CD3 complex. The relative subtle TCR expression defect seen in the absence of CD3γ mirrors the relatively subtle clinical phenotype of patients with γ-deficiency compared to the severe disease of patients with δ-deficiency(17)

We have previously demonstrated that amino acid changes in the framework of the variable α and β domains can enhance TCR expression without changing antigen specificity(7). Here, we have demonstrated that combining TCR V-region engineering and CD3ζ gene transfer can further increase TCR expression and antigen-specific T cell function. Compared to T cells transduced with wild type TCRs, transduction with V-region engineered TCR and CD3ζ enhanced antigen-specific T cell responses at least 3-fold at all peptide concentrations tested. While the benefit of adding CD3ζ to the V-region engineered CMV-TCR was small, the performance of the engineered HA1-m2 and HA1-m7 TCR was doubled by the presence of ζ. The data in this report support the inclusion of the CD3ζ gene in vectors used for TCR gene therapy in humans.

Recent publications have shown the benefits of disrupting endogenous TCR genes in human T cells engineered to express chimeric antigen receptors (CARs) or TCRs. The insertion of CARs into the TCRα gene locus has improved the ability of T cells to protect against tumour growth in murine models(18). In this setting, CAR expression was regulated by the endogenous TCRα promoter, which resulted in lower levels of expression compared to CAR expression from a retroviral promotor, and functionally correlated with reduced constitutive CAR signalling and reduced T cell exhaustion. The benefit of knocking out endogenous TCRα and β genes and inserting an exogenous TCRα/β construct into the TCRα locus under the control of the endogenous promoter has also been demonstrated recently(19). In this setting, removing endogenous TCR α and β chains prevented any mispairing with the introduced TCR chains. However, the surface expression level of the introduced TCR was not determined by the TCRα promoter but remain instead under the control of the endogenous CD3ζ chain. Hence, it remains to be explored whether expression of introduced TCRs from the endogenous TCRα promotor provides T cells with functional benefits over TCR expression driven by promoters of lenti and retroviral vectors, as in each case the regulation of surface expression remains under the control of endogenous CD3ζ.

## Author Contributions

AD designed and conducted experiments, analysed data and wrote the paper. ST and AH designed experiments and wrote the paper. HS initiated the study, designed experiments, analysed data and wrote the paper. All authors contributed to the article and approved the submitted version.

## REFERENCES

1. Brenner MB, McLean J, Dialynas DP, Strominger JL, Smith JA, Owenll FL, Seidman JG, Ip S, Rosen F, Kranger tt MS. Identification of a putative second T-cell receptor.

2. Call ME, Pyrdol J, Wiedmann M, Wucherpfennig KW. The Organizing Principle in the Formation of the T Cell Receptor-CD3 Complex tion of each polar residue or the resulting arrangement. The working hypothesis has been that these basic and acidic residues form pairwise interactions in the mem. (2002). 967–979 p. http://www.cell.com/

3. Call ME, Wucherpfennig KW. The T cell receptor: Critical role of the membrane environment in receptor assembly and function. Annu Rev Immunol (2005) 23:101– 125. doi: 10.1146/annurev.immunol.23.021704.115625

4. Call ME, Wucherpfennig KW. The T cell receptor: Critical role of the membrane environment in receptor assembly and function. Annu Rev Immunol (2005) 23:101– 125. doi: 10.1146/annurev.immunol.23.021704.115625

5. Ahmadi M, King JW, Xue SA, Voisine C, Holler A, Wright GP, Waxman J, Morris E, Stauss HJ. CD3 limits the efficacy of TCR gene therapy in vivo. Blood (2011) doi: 10.1182/blood-2011-04-346338

6. Degirmencay A, Thomas S, Mohammed F, Willcox BE, Stauss HJ. Modifications outside CDR1, 2 and 3 of the TCR variable β domain increase TCR expression and antigen-specific function. Front Immunol (2023) 14: doi: 10.3389/fimmu.2023.1148890

7. Thomas S, Mohammed F, Reijmers RM, Woolston A, Stauss T, Kennedy A, Stirling D, Holler A, Green L, Jones D, et al. Framework engineering to produce dominant T cell receptors with enhanced antigen-specific function. Nat Commun (2019) doi: 10.1038/s41467-019-12441-w

8. Sušac L, Vuong MT, Thomas C, von Bülow S, O’Brien-Ball C, Santos AM, Fernandes RA, Hummer G, Tampé R, Davis SJ. Structure of a fully assembled tumor-specific T cell receptor ligated by pMHC. Cell (2022) 185:3201–3213.e19. doi: 10.1016/j.cell.2022.07.010

9. Dong D, Zheng L, Lin J, Zhang B, Zhu Y, Li N, Xie S, Wang Y, Gao N, Huang Z. Structural basis of assembly of the human T cell receptor–CD3 complex. Nature (2019)

10. Degirmencay A, Thomas S, Mohammed F, Willcox BE, Stauss HJ. Modifications outside CDR1, 2 and 3 of the TCR variable β domain increase TCR expression and antigen-specific function. Front Immunol (2023) 14: doi: 10.3389/fimmu.2023.1148890

11. Baniyash M. TCR ?-chain downregulation: Curtailing an excessive inflammatory immune response. Nat Rev Immunol (2004) 4:675–687. doi: 10.1038/nri1434

12. Kulkarni DP, Wadia PP, Pradhan TN, Pathak AK, Chiplunkar S V. Mechanisms involved in the down-regulation of TCR ? chain in tumor versus peripheral blood of oral cancer patients. Int J Cancer (2009) 124:1605–1613. doi: 10.1002/ijc.24137

13. Region U, Chowdhury B, Krishnan S, Tsokos CG, Robertson JW, Fisher CU, Nambiar MP, Tsokos GC. Stability and Translation of TCR. (2006)1–11. papers3://publication/uuid/25EB5983-AF42-4B52-829C-375B1FB85BF9

14. Kono K, Salazar-Onfray F, Petersson M, Hansson J, Masucci G, Wasserman K, Nakazawa T, Anderson P, Kiessling R. Hydrogen peroxide secreted by tumor-derived macrophages down-modulates signal-transducing zeta molecules and inhibits tumor-specific T cell-and natural killer cell-mediated cytotoxicity. Eur J Immunol (1996) 26:1308–1313. doi: 10.1002/eji.1830260620

15. Rabinowich H, Reichert TE, Kashii Y, Gastman BR, Bell MC, Whiteside TL. Lymphocyte apoptosis induced by Fas ligand-expressing ovarian carcinoma cells: Implications for altered expression of T cell receptor in tumor-associated lymphocytes. Journal of Clinical Investigation (1998) 101:2579–2588. doi: 10.1172/JCI1518

16. Aguinaga-Barrilero A, Castro-Sánchez P, Juárez I, Gutiérrez-Calvo A, Rodríguez-Pérez N, Lopez A, Gómez R, Martin-Villa JM. Defects at the posttranscriptional level account for the low TCR? chain expression detected in gastric cancer independently of caspase-3 activity. J Immunol Res (2020) 2020: doi: 10.1155/2020/1039458

17. Recio MJ, Moreno-Pelayo MA, Kiliç SS, Guardo AC, Sanal O, Allende LM, Pérez-Flores V, Mencía A, Modamio-Høybjør S, Seoane E, et al. Differential Biological Role of CD3 Chains Revealed by Human Immunodeficiencies. The Journal of Immunology (2007) 178:2556–2564. doi: 10.4049/jimmunol.178.4.2556

18. Eyquem J, Mansilla-Soto J, Giavridis T, Van Der Stegen SJC, Hamieh M, Cunanan KM, Odak A, Gönen M, Sadelain M. Targeting a CAR to the TRAC locus with CRISPR/Cas9 enhances tumour rejection. Nature (2017) 543:113–117. doi: 10.1038/nature21405

19. Schober K, Müller TR, Gökmen F, Grassmann S, Effenberger M, Poltorak M, Stemberger C, Schumann K, Roth TL, Marson A, et al. Orthotopic replacement of T-cell receptor α- and β-chains with preservation of near-physiological T-cell function. Nat Biomed Eng (2019) doi: 10.1038/s41551-019-0409-0

